# Punctuated loci on chromosome IV determine natural variation in Orsay virus susceptibility of *Caenorhabditis elegans* strains Bristol N2 and Hawaiian CB4856

**DOI:** 10.1101/2020.10.30.361683

**Authors:** Mark G. Sterken, Lisa van Sluijs, Yiru A. Wang, Wannisa Ritmahan, Mitra L. Gultom, Joost A.G. Riksen, Rita J.M. Volkers, L. Basten Snoek, Gorben P. Pijlman, Jan E. Kammenga

## Abstract

Host-pathogen interactions play a major role in evolutionary selection and shape natural genetic variation. The genetically distinct *Caenorhabditis elegans* strains, Bristol N2 and Hawaiian CB4856, are differentially susceptible to the Orsay virus (OrV). Here we report the dissection of the genetic architecture of susceptibility to OrV infection. We compare OrV infection in the relatively resistant wild-type CB4856 strain to the more susceptible canonical N2 strain. To gain insight into the genetic architecture of viral susceptibility, 52 fully sequenced recombinant inbred lines (CB4856 x N2 RILs) were exposed to OrV. This led to the identification of two loci on chromosome IV associated with OrV resistance. To verify the two loci and gain additional insight into the genetic architecture controlling virus infection, introgression lines (ILs) that together cover chromosome IV, were exposed to OrV. Of the 27 ILs used, 17 had an CB4856 introgression in an N2 background and 10 had an N2 introgression in a CB4856 background. Infection of the ILs confirmed and fine-mapped the locus underlying variation in OrV susceptibility and we found that a single nucleotide polymorphism in *cul-6* contributes to the difference in OrV susceptibility between N2 and CB4856. An allele swap experiment showed the strain CB4856 became more susceptible by having an N2 *cul-6* allele, although having the CB4856 *cul-6* allele did not increase resistance in N2. Additionally, we found that multiple strains with non-overlapping introgressions showed a distinct infection phenotype from the parental strain, indicating that there are punctuated locations on chromosome IV determining OrV susceptibility. Thus, our findings reveal the genetic complexity of OrV susceptibility in *C. elegans* and suggest that viral susceptibility is governed by multiple genes.

## Introduction

Genetic variation plays a major role in the arms race between pathogen and host [1–3]. The interaction between host genetic background and pathogen can shape natural variation by imposing a strong selection regime on the affected population. Host genetic variation plays a role in ongoing viral outbreaks as illustrated by studies that correlate outcome of infection with Hepatitis, HIV, Zika, Ebola and SARS-CoV-2 to the host’s genetic background [4–9]. Studying host-virus interactions in model systems can uncover genetic networks determining viral susceptibility [10].

The nematode *Caenorhabditis elegans* encounters a variety of pathogens in its natural habitat, including bacteria, microsporidia, oomycetes, and fungi [11]. So far, only one virus has been discovered that naturally infects *C. elegans*: the Orsay virus (OrV) [12]. In the laboratory this pathogen can be easily maintained and used to study host-virus interactions [12]. Host-virus interaction studies focusing on the effect of host genetic variation are facilitated by the androdiecious mode of replication by which *C. elegans* reproduces. This makes *C. elegans* a powerful model to investigate the effect of host genetic variation as populations can be both inbred and outcrossed.

Three cellular pathways are used by *C. elegans* to defend itself against viral infections. First, the RNAi response is a highly adaptive and diverse pathway that plays a role in many processes in an organism, for example in development and antiviral responses in invertebrates [13,14]. In OrV infection, it recognizes the double stranded RNA replication intermediate, which ultimately leads to the production of small interfering RNAs (siRNAs) that target the viral RNA for degradation [15–19]. Mutants defective for various genes in the RNAi pathway display higher viral susceptibility upon infection [15,17–19]. Second, the OrV can be targeted by a distinct mechanism known as viral uridylation [20]. Uridylation, like RNAi, leads to degradation of viral RNAs although both antiviral defenses function independently of one another. Third, the Intracellular Pathogen Response (IPR) is involved in defense against viral, fungal and microsporidian infections. The IPR is regulated by the gene pair *pals-22* and *pals-25* that balance the nematode’s physiological programs between growth and immunity [21]. Infections are counteracted by upregulating a range of 80 IPR genes that reduce proteotoxic stress [21–23]. For most IPR genes, their biochemical function is currently unknown, but IPR gene *cul-6* functions in the E3 ubiquitin ligase complex and protects against viral and microsporidian infection [23,24]. Furthermore, the gene *drh-1* (encoding a RIG-I like protein) mediates the IPR response specifically upon OrV infection connecting IPR and RNAi pathways which both depend on this gene [25].

Natural variation influences the susceptibility to OrV infections. Initially, it was observed that the natural *C. elegans* strain JU1580 is more susceptible to infection with OrV than the reference strain Bristol N2 [18]. This difference has been linked to a natural polymorphism in *drh-1* affecting the anti-viral RNAi response [15]. In addition to the natural variation in the RNAi response, genetic variation also determines the Intracellular Pathogen Response (IPR) against OrV infection. The Hawaiian strain CB4856 had higher (basal) expression of multiple IPR genes than N2, potentially resulting in higher resistance to OrV infection observed in CB4856 [26]. However, the genetic and transcriptional networks leading to this difference have not been uncovered.

The CB4856 and N2 strain are very polymorphic, with more than 400,000 polymorphisms, including insertions/deletions and single nucleotide variants [27,28]. Over the last decade, both strains have been jointly used in many quantitative genetics studies in *C. elegans*, focused on traits like: aging, stress tolerance and pathogen avoidance [29–33]. Most of these studies have been conducted on one of the two available recombinant inbred line (RIL) panels [34,35] or on the introgression line (IL) population which contains fragments of CB4856 in a background of N2 [30].

Here we set out to investigate genetic loci involved in the phenotypic differences between the Bristol N2 strain and the Hawaii CB4856 in response to OrV infection. Viral replication was characterized in N2 and CB4856 in a stage- and incubation time-dependent manner. Subsequently, we used inbred panels constructed from these strains to identify possible causal loci underlying the difference in viral susceptibility. We exposed a panel of 52 RILs to OrV and measured the viral load. We identified two QTL associated with differences in viral load on chromosome IV. Following-up, using a panel of 27 IL strains together covering the QTL location on chromosome IV led to the identification of 34 candidate genes involved in antiviral immunity. One of these candidate genes, the IPR gene *cul-6*, was tested for its role in OrV infection in the strains N2 and CB4856.

## Material and methods

### C. elegans strains

*C. elegans* strains Bristol N2 and Hawaii CB4856 were used and strains derived by crossing these two wild-type strains. In this paper 52 Recombinant Inbred Lines (RILs), 17 Introgression Lines with an N2-background (IL_N2_) and 10 Introgression Lines with a CB4856 background (IL_CB4856_) covering chromosome IV were used. IL strains are described in Supplementary Table S1 and RIL strains were described previously [35]. All these genotypes have been confirmed by whole-genome sequencing on the Illumina HiSeq 2500 platform as described by previously [36]. The strains PHX1169 *cul-6(syb1169)* and PHX1170 *cul-6(syb1170)*, containing the *cul-6* CB4856 allele in a N2 background and the N2 *cul-6* allele in a CB4856 background respectively, have been created by CRISPR-Cas9 by SunyBiotech (http://www.sunybiotech.com) (Supplementary Text S1). These genotypes have been confirmed by PCR sequencing.

### C. elegans culturing

The nematodes were kept at 12°C in-between experiments on 6 cm NGM plates seeded with *E. coli* OP50. Bleaching was used to synchronize populations and to remove bacterial or fungal contaminations [37]. Before experiments, a population without males was created by picking single worms in the L1/L2 stage and transferring hermaphrodite populations to fresh 9 cm NGM plates. New experiments were started by bleaching an egg-laying population grown at 20°C.

### Orsay virus stock preparation

Orsay virus stocks were generated by isolating OrV from a persistently infected JU1580 culture as previously described [17,18]. In short, JU1580 populations were grown on 100 9 cm NGM plates [37] containing twice the usual amount of agar to prevent the nematodes from burying into the agar (34 g/L). The nematodes were collected by washing the animals off the plate with M9 buffer [37] and flash freezing the suspension in liquid nitrogen. After defrosting on ice, the supernatant was collected and passed through a 0.2 μm filter. Specific infectivity of the virus stock was tested by serial dilution infections in *C. elegans* JU1580 [17].

### Infection experiments

The infection assay was conducted as described previously [17]. Populations were synchronized (t=0 hours) and grown at 20°C on 9 cm NGM plates. Infections were performed on animals in the L1 (22 hours post bleaching) or L2 (26 or 28 hours post bleaching as indicated in the text) stage. Prior to the infection, the nematodes were washed off the plate with M9 buffer and pelleted by centrifugation. The supernatant was removed, and the nematodes were exposed to OrV in liquid for 1 hour. The worms were washed 3 times with M9 and placed on a fresh 9 cm NGM plate.

For the replication kinetics experiments on N2 and CB4856, the animals were harvested 2-35 hours post infection. This experiment was conducted in 8 independent biological replicates, each evenly covering the time-series. For the viral load experiments on the RIL and IL panels and the *cul-6* allele swap strains, the animals were harvested 30 hours post infection. The experiment in the RIL panel was conducted on 3 independent biological replicates. The experiment in the IL panel was conducted on at least 5 independent biological replicates. The experiment using the *cul-6* allele swap strains was conducted on 21 independent biological replicates.

### RNA isolation

The RNA was isolated using a Maxwell® 16 AS2000 instrument with a Maxwell® 16 LEV simply RNA Tissue Kit (both Promega) following the recommended protocol, except the addition of 10 mg of proteinase K during the lysis step. The lysate was incubated in a Thermomixer (Eppendorf) for 10 minutes at 65°C at 1,000 rpm. After isolation the quality and quantity of the RNA was determined via NanoDrop (Thermo Scientific).

### cDNA preparation and qPCR

cDNA was synthesized using the GoScript Reverse Transcriptase kit (Promega) following the recommended protocol with random hexanucleotides (Thermo Scientific) and 1 μg of total RNA as starting material. The cDNA was quantified by qPCR (MyIQ, Biorad) using Absolute QPCR SYBR Green Fluorescein Mixes (Thermo Scientific) or iQ SYBR Green Supermix (Biorad) following the recommended protocol. The samples were quantified using the primers described by [17].

The qPCR data was processed using R (version 4.0.2), as described before [17]. In short, before normalization, the qPCR measurements were transformed by

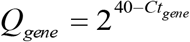

where Q_gene_ is the expression of the gene and Ct_gene_ is the measured Ct value of the gene. The viral expression was normalized by the two reference genes, using the formula

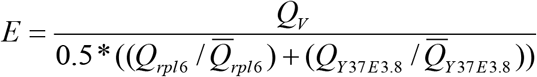

where E is the normalized viral load, Q_V_ is the expression of the viral RNA and Q_*rpl6*_ and Q_Y37E3.8_ are the expression of reference genes *rpl-6* and Y37E3.8 respectively. Viral load data presented here was batch correct for the batch effect caused by the different virus stock used by correcting for the average viral loads of N2 and CB4856 (excluding unsuccessful infections) as these two strains were taken along in every experiment.

From the replicate measurements in the RIL panel, several traits could be derived for QTL mapping over the RIL population. The following parameters were derived including all measurements: mean viral load, median viral load, and minimum viral load. We excluded the unsuccessful infections (as these could arise due to technical failures) unless indicated otherwise.

### Quantitative trait locus mapping RIL population

Single locus QTL mapping was done using a linear model in R (version 4.0.2) to explain viral load and derived traits over the markers by

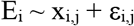

where E is the viral load of RIL i (1, 2, …, 52) and x is the marker of RIL i at location j (a set of 1152 sequenced markers was used (Supplementary Table S2) [27]. For E the outcome of each replicate of the experiment was averaged over the three biological replicates.

For the QTL mapping the statistical threshold was determined via a permutation analysis, where the values measured for E were randomly distributed over the genotypes. The same model as for the mapping was used and this analysis was repeated 1,000 times. The 950th highest p-value was taken as the p-value threshold for a false discovery rate of 0.05.

### Heritability and variance calculations

The narrow-sense heritability’s (*h^2^*) were calculated per investigated trait by REML [14,38,39] using the package ‘heritability’ in R (version 4.0.2). Significance was determined via 1,000 permutations where the values measured for E were randomly distributed over the genotypes.

The variation of viral loads explained by a QTL peak (V_Explained_) was calculated by

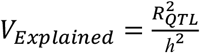

where R^2^_QTL_ is the determination coefficient from fitting the peak marker to the trait as calculated by a linear model and *h^2^* is the narrow-sense heritability of the trait.

### Introgression line analysis

The viral loads obtained for the introgression lines were analyzed individually against N2 and CB4856 via a two-sided t-test assuming unequal variance in R (version 4.0.2). Experiments where no virus was detected were excluded from the analysis. Moreover, we performed linkage mapping for the two IL panels separately using a linear model to explain viral load over the markers by

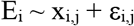

where E is the viral load of IL i (1, 2, …, 10 or 17) and x is the marker of IL i at location j (a set of 1152 sequenced markers was used (Supplementary S2) Each IL was compared against the respective parental strain (N2 or CB4856). For E the outcome of each replicate of the experiment was averaged over the biological replicates. A significance threshold was drawn at −log_10_(p) > 3.5 for analysis of the data.

### Allele swap analysis

Because we observed a high level of variance in the viral loads in N2 and CB4856 and the effect size of the QTL_IV:12.41-12.89_ was small we used a high level of replication for the allele swap experiments by performing 21 biologically independent infections using three different virus stocks. Unsuccessful infections were excluded from the analysis and the batch corrected viral load data (based on virus stock as described above) was subsequently checked for outliers. Outliers were defined by 1.5 times the interquartile range plus or minus the third or first quartile respectively. After removal of the outliers (7% of the measurements), a t-test assuming unequal variances was performed to test for differences in viral load.

### Protein structure analysis

Protein sequences from the human CUL1 (NCBI Reference Sequence: NP_003583.2), *Saccharomyces cerevisiae* CDC53 (GenBank: CAA98702.1), *Drosophila melanogaster* CUL-1 (GenBank: AAD33676.1) and *C. elegans* CUL-1 (GenBank: AAC47120.1), CUL-6 N2 allelic variant (GenBank: CAB01230.1) and CUL-6 CB4856 allelic variant were aligned using ClustalX (version 2.1) with the default settings [40]. A structural model for the N2 and CB4856 allelic variant was predicted using the human CUL1 protein structure as a template in the SWISS-MODEL ExPASy web server. The default search parameters were used, based on the SWISS-MODEL template library (version 14/01/2015) and the protein data bank (version 09/01/2015) [41–46]. The obtained models for N2 and CB4856 CUL-6 were compared in SwissPDBViewer (v. 4.1.0) [47].

### Script availability

Custom scripts written in R (version 4.0.2) are openly available at https://git.wur.nl/published_papers/sterken_sluijs_2020.

## Results

### CB4856 displays resistance to OrV infection

The infection kinetics of OrV were investigated in the two wild-type strains N2 and CB4856. Infection kinetics were investigated by infecting both strains at an age of 26 hours (L2 stage) and measuring the viral load over 2-35 hours post infection (in 28-61 hour old animals) (Figure 1A). N2 developed a higher maximum viral load than CB4856 in this time period (Figure 1B). The infection developed via a clear lag-phase in N2 during the first 12 hours, whereas large variation in viral loads was observed in the initial infection phase for CB4856 (Figure 1B). In this time series experiment, a significant amount of the variance was explained by the different genotypes (ANOVA, p < 1·10^−4^). We found that for some infected CB4856 populations the infection developed via a similar pattern (but to lower viral load) compared to N2, however in other experiments the infection did not develop beyond levels reached in the lag-phase of the infection. Consequently, CB4856 populations that were 38h or older showed either similar viral loads to populations that were younger (and thus shorter infected) or viral loads that reached the maximum viral load for CB4856 (Figure 1C). On the other hand, N2 populations all reached higher viral loads after the lag-phase of infection was passed (Figure 1C). Therefore, the time passed since infection also explained variation in viral load (ANOVA, p < 1·10^−6^). Next to this, we observed that infection tended to be more often established in N2 (76% success rate) than in CB4856 (61% success rate) (chi-square test, p = 0.093). Constant exposure to OrV for four days resulted in similar viral loads between N2 and CB4856 [15,26], thus suggesting that multiple rounds of viral replication are necessary to fully infect CB4856 populations. Together, these observations show that CB4856 develops a lower viral load and can suppress a beginning infection better than N2.

**Figure 1.**
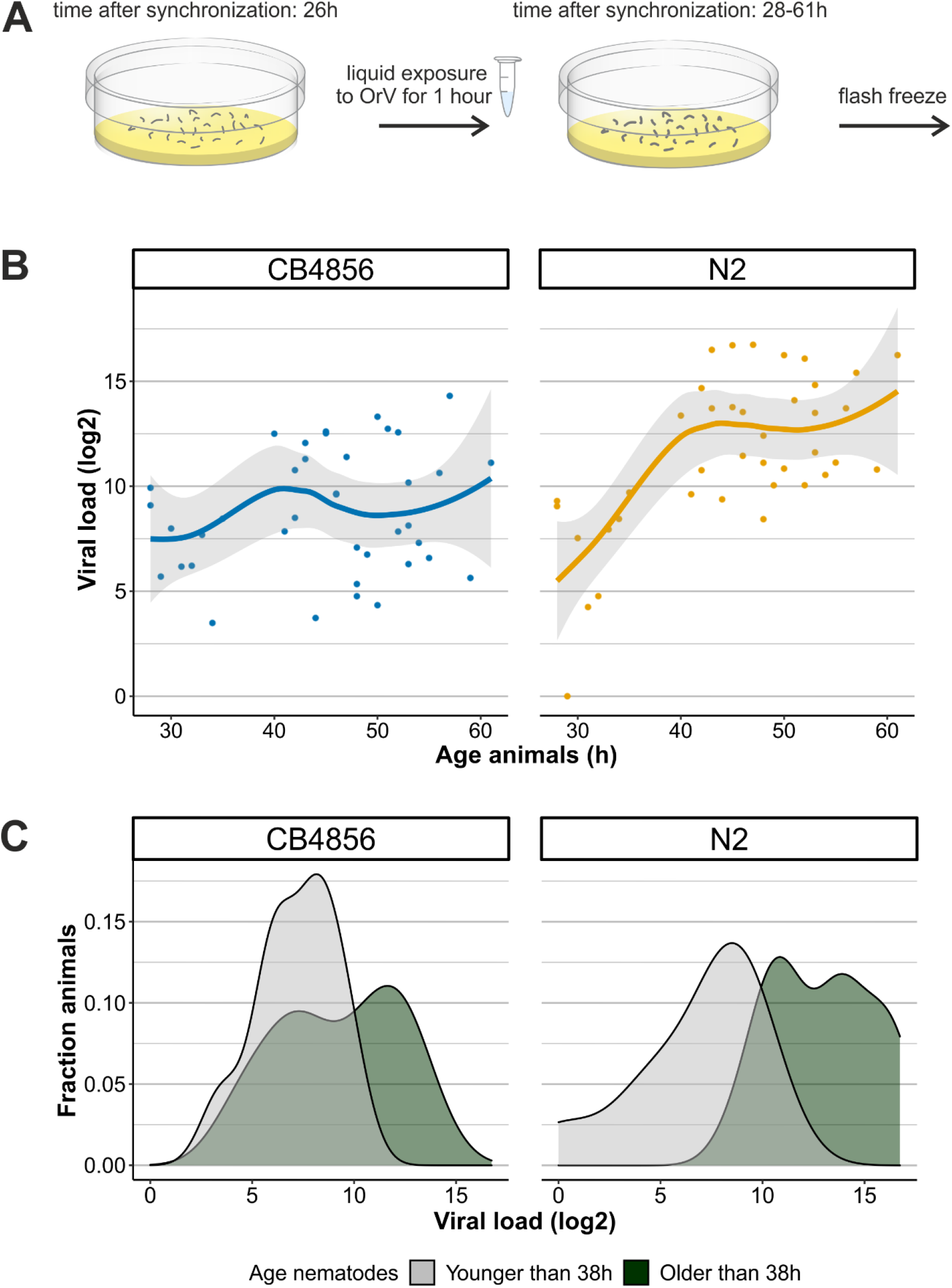
Kinetics of OrV infection in N2 and CB4856. A) Nematodes are infected by the OrV in liquid at the age of 26h hours (as in [17]) before samples were washed of the plate 2-35h later and collected for viral load quantification. B) Development of OrV infection over time in N2 and CB4856 over a course of 35h. The points depict the observed viral loads, the line represents the smoothed conditional mean and the grey shading shows the confidence interval for the mean. C) A density plot of the viral load measurements over time, divided in two groups: early infection (up to 12 hours post infection, grey) and late infection (after 12 hours post infection, green).

A reason for the difference in viral load between CB4856 and N2 could be a stage dependent difference in resistance as was found for CB4856 nematodes, which are resistant to infection by the microsporidian *Nematocida parisii*, but only in the L1 stage [48]. Moreover, *N. parisii* shares its cellular tropism with OrV and both pathogens induce the same transcriptional response: the Intracellular Pathogen Response (IPR) [21,22,49]. Therefore, we also tested if L1 CB4856 could exhibit even higher resistance to OrV infection than the L2 animals we have infected before. Infection was compared in first (22-hour old) and second (28-hour old) larval stage animals (Figure S1A). N2 animals were infected in parallel for reference and the infection could develop for 30 hours after infection. We found for both genotypes that the viral loads were highly comparable between L1 and L2 infected nematodes (Figure S1B). Thus, the relative resistance of CB4856 towards the OrV is not stage-dependent, in contrast to resistance to the microsporidian *N. parisii*.

### A locus on chromosome IV links to resistance against OrV

To find the causal genetic loci underlying the different viral loads between N2 and CB4856 in viral load, recombinant inbred lines (RILs) constructed from a cross between these strains were infected with OrV (Figure S2A) [27,35]. The RILs were infected in the L2 stage (at the age of 26 hours) and the infection was continued for 30 hours, after which the viral load was measured. The viral loads of the RILs followed a pattern of transgressive segregation, indicating that multiple genetic loci contribute to viral susceptibility (Figure 2A). We found a narrow-sense heritability (*h^2^,* the fraction of trait variation explained by genotype) at 0.40 for the mean viral load (excluding populations that were not successfully infected) meaning that 40% of this phenotype can be explained by additive genetic variance. Linkage analysis for this trait identified a QTL on chromosome IV between 12.5 and 15.1Mb (Figure 2B) (R^2^ = 0.37). Besides performing a linkage analysis for the mean viral load of successfully infected populations, linkage analysis was performed for A) the median viral load (excluding unsuccessfully infected populations), B) the overall mean viral load (including unsuccessfully infected populations) and C) the minimum viral load observed for a strain (Figure S2A-C). Correlation analysis of the minimum viral load pointed towards an additional QTL location between 2.6 and 2.8Mb on chromosome IV (Figure S2C). Thus, this QTL location could link to the success of infection, whereas the QTL location on the right side of chromosome IV was linked to the height of the viral load measured. Therefore, each locus may influence another biological aspect of OrV infection.

**Figure 2.**
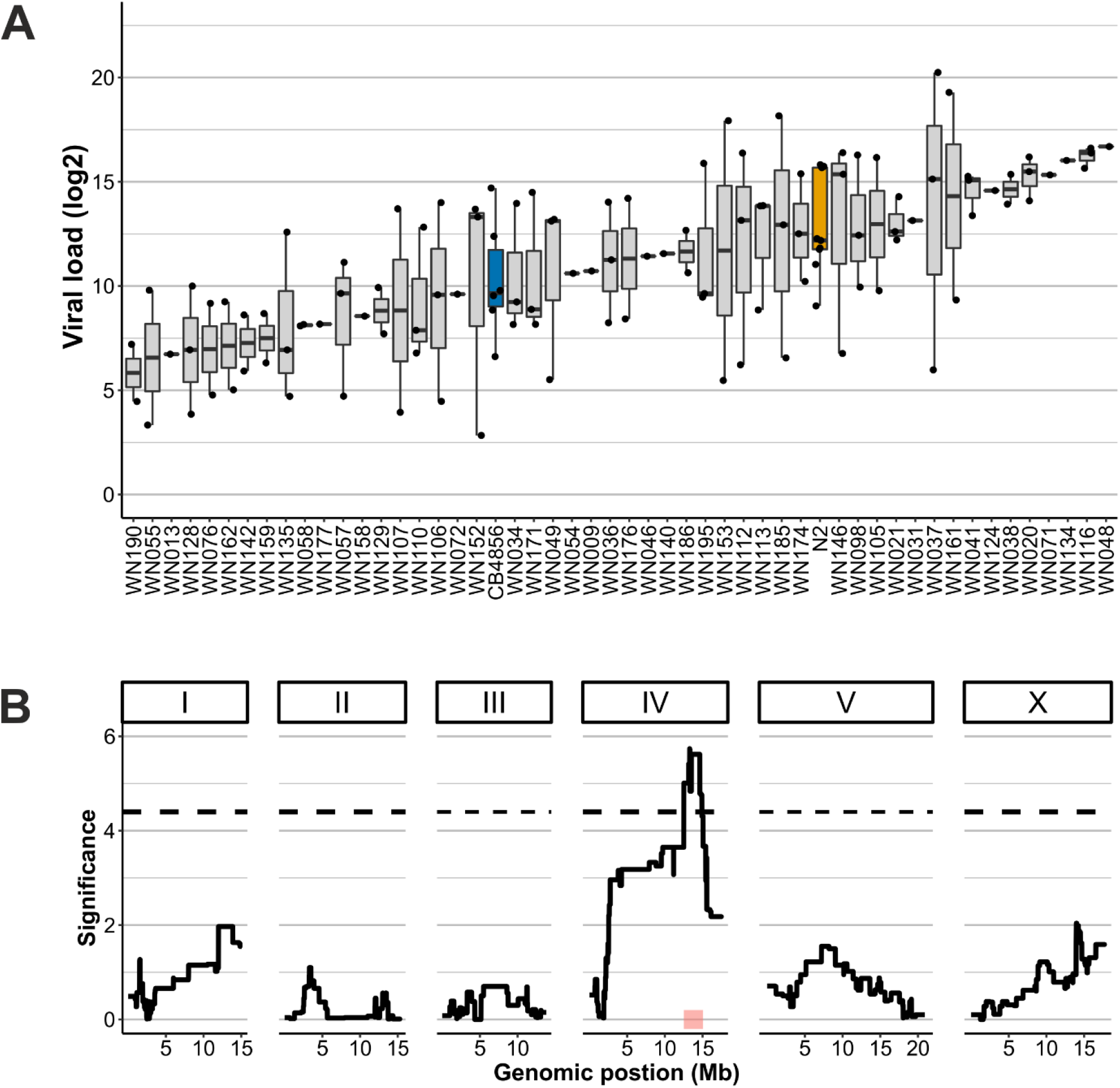
OrV infections in a Recombinant Inbred Line panel with parental strains N2 (orange) and CB4856 (blue) and QTL mapping. A) Transgression plot of the viral loads of 52 RIL strains used for the infection assays Each dot represents a biological replicate of a viral load measured after infection. All RIL strains were infected three times, but samples that lacked viral replication (viral load = 0) are not shown. B) The QTL profile for mean viral load (excluding unsuccessful infections). The significant QTL peak is found at the end of chromosome IV at 13.3Mb (1.5 LOD-drop interval from 12.5-15.1Mb) as indicated by the red square.

### Verification of the QTL locus by introgression lines

To experimentally verify the QTL involved in the viral susceptibility difference between N2 and CB4856 introgression lines (ILs) were infected. ILs contain small fragments of one strain in the genetic background of another strain [30]. ILs that together cover chromosome IV were used and their viral loads were measured after infection. We used 10 ILs with a N2 fragment in the CB4856 background (IL_CB4856_) and 17 ILs with a CB4856 fragment in the N2 background (IL_N2_; Figure S2B). Of the 27 infected ILs, nine had a different viral load than the parental strain, demonstrating that presence of the introgression alters the viral susceptibility compared to the parent. We found that the IL_CB4856_ strains WN352, WN353, and WN354 showed a phenotype distinct from the parental CB4856 strain (two-sided t-test, p < 0.05). These strains carry introgressions that together overlapp the right QTL peak at 12.41-12.89Mb. In agreement, two IL_N2_ strains covering this QTL were more resistant than N2 (WN252, WN254) (Figure 3A), but contrary three strains with the CB4856 fragment in the N2 background covering the same location did not show a lower viral load than the N2 strain (WN258, WN259, and WN261). In addition, IL_N2_ strain WN263 with an introgression from 14.87-17.49Mb had a lower susceptibility than N2. These results indicate that there are multiple loci underlying the susceptibility difference between N2 and CB4856 that are likely to interact together.

**Figure 3.**
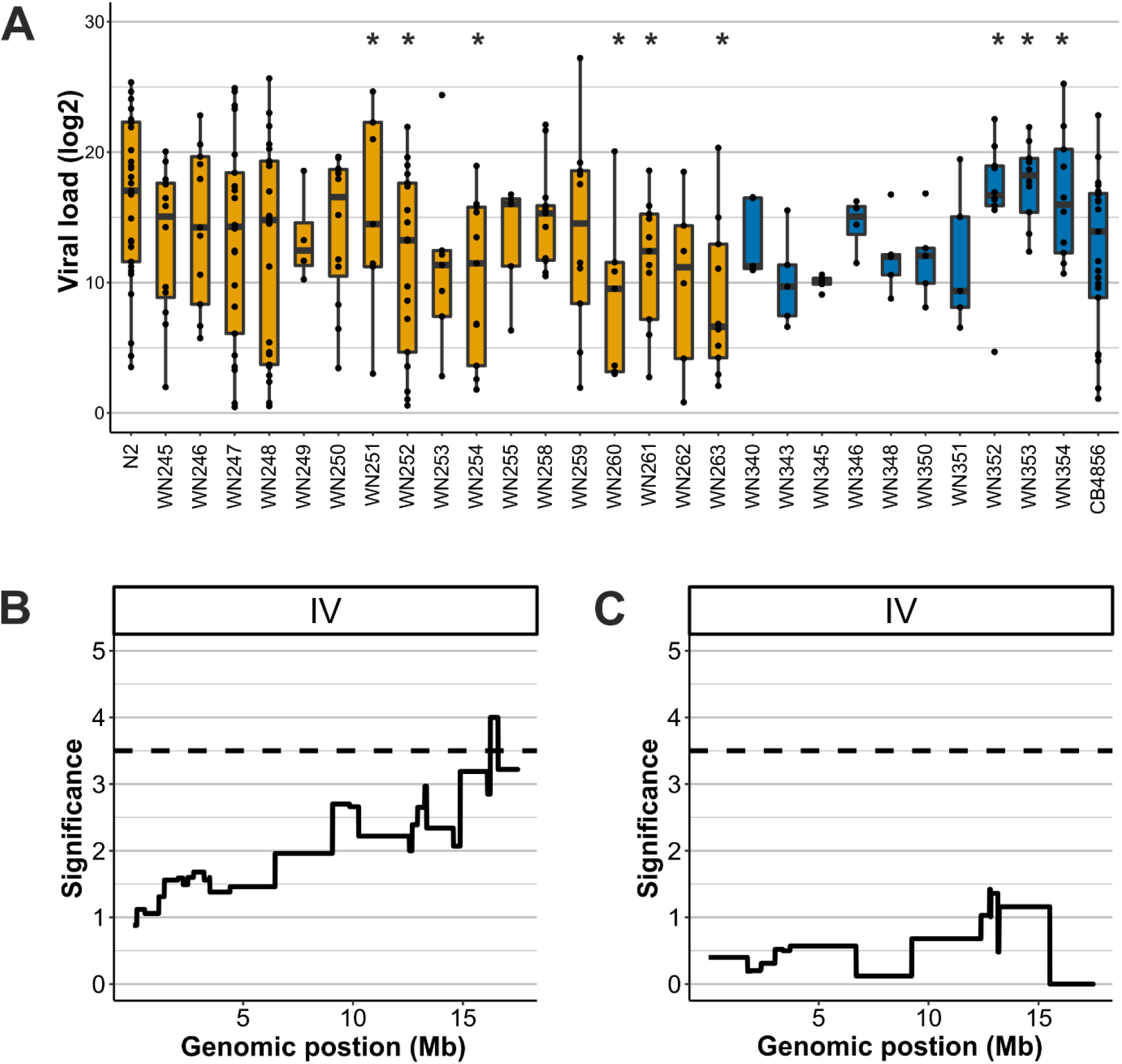
OrV infections in two Introgression Line panels with parental strains N2 and CB4856 and QTL mapping. A) The viral loads of N2, CB4856 and 27 IL strains used for the infection assays. Of these, 17 strains have a CB4856 introgression in a N2 background (orange filled boxplots) and 10 have a N2 introgression in a CB4856 background (blue filled boxplots). An asterisk indicates a significant difference from its parental genetic background (p < 0.05, t-test). Each dot represents a biological replicate. Samples that lacked viral replication (viral load = 0) are not shown. B) Linkage mapping profile for mean viral load (excluding unsuccessful infections) for the IL_N2_ background panel. A significant peak is found on the right side of chromosome IV. C) Linkage mapping profile for mean viral load (excluding unsuccessful infections) for the IL_CB4856_ background panel.

Linkage analysis on the IL_N2_ panel showed the highest correlations for mean viral load and genetic background on the right side of chromosome IV with a QTL peak around 16Mb (Figure 3B), whereas the IL_CB4856_ panel mapping did not show an effect of the introgression (Figure 3C). The resolution for QTL mapping in the ILs is relatively low compared to QTL mapping in the RILs, because of fewer genetic breakpoints in the population. Therefore, the peak mapped in the ILs could rely on the same genetic variation as the QTL peak mapping in the RIL panel that estimated a QTL between 12.5-15.1Mb. Furthermore, we could not find a simple conformation for the 2.6-2.8Mb QTL, although two (WN251 and WN252) out of fourteen strains covering this locus indicate an effect in certain genetic backgrounds. .

### Candidate causal genes underlying different viral susceptibility between N2 and CB4856

Linkage analysis in both RILs and ILs indicated that viral susceptibility differences between N2 and CB4856 were governed by multiple loci. So, we set out to see if we could identify polymorphic genes that determine the difference in viral susceptibility. We focus on the 12.41-12.89Mb region on chromosome IV, because this region was mapped in the RIL panel and supported by analysis of the ILs. This region contains 34 polymorphic genes of which 25 contain a non-synonymous change in the coding sequence (Supplementary Table S3). The candidate genes in this region have diverse functions, including genes with a known immune function against bacterial or viral infection. One of these is *cul-6*, which is regulated by the IPR. A knockdown of *cul-6* increases the susceptibility to OrV in N2 nematodes [21,22,24]. The CB4856 allele of *cul-6* gene contains a single nucleotide polymorphism in the 428^th^ amino acid, physically close to the RBX-1 binding site, where a negatively charged glutamic acid is found in N2 and a positively charged lysine in CB4856 (Figure 4A). The amino acid lysine at this position has been highly conserved from yeast to humans in the closely related CDC53 and CUL-1 proteins (amino acid conservation between *C. elegans* CUL-1 and CUL-6 is 47%) (Figure 4B) [50].

**Figure 4.**
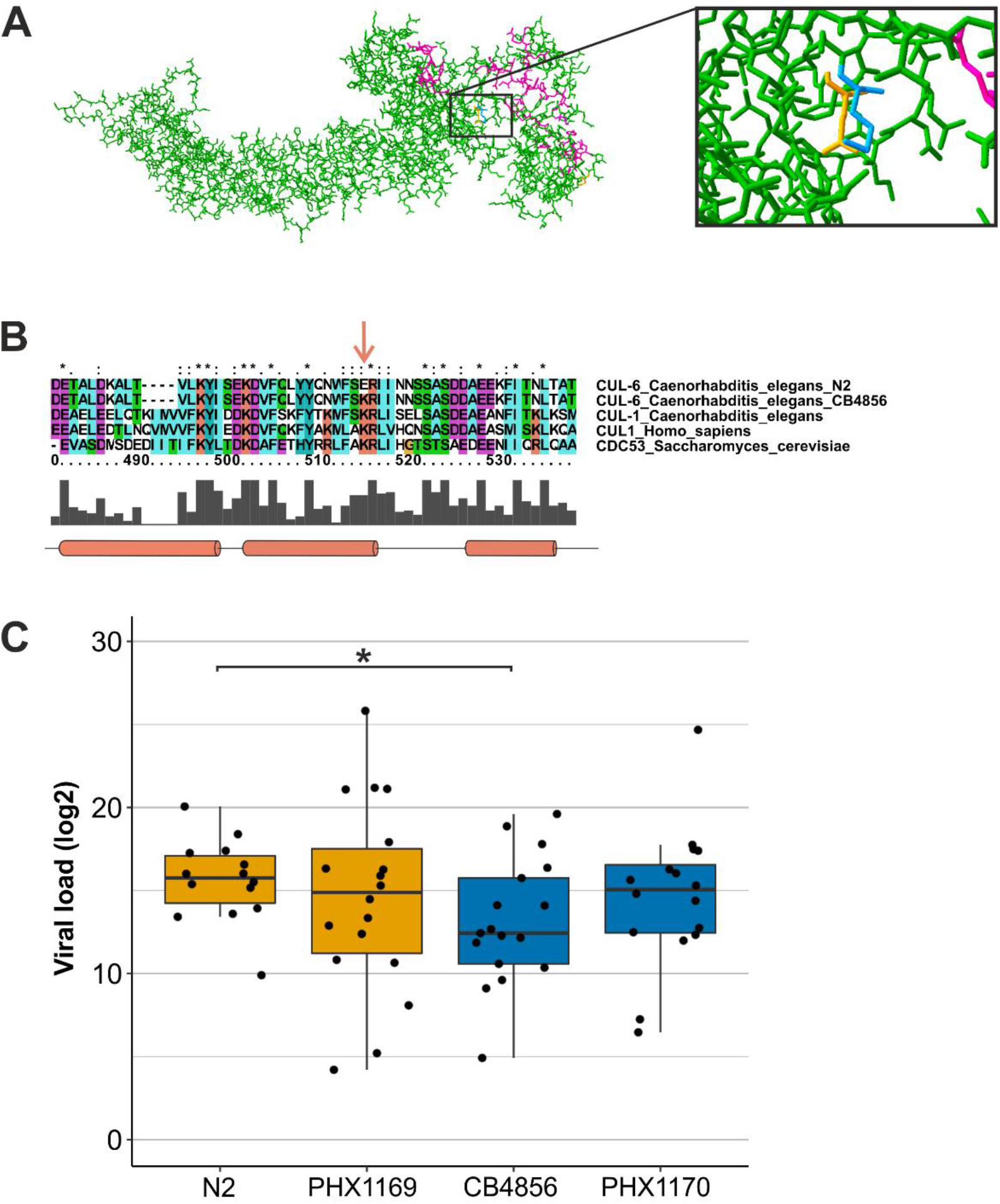
The cul-6 gene in CB4856 and N2 and its effect on viral susceptibility. A) Structure prediction of C. elegans CUL-6. The lysine present in the CB4856 allelic variant is shown is blue, the glutamic acid present in the N2 allelic variant in orange. The RBX-1 binding domain is shown in purple. B) Part of the sequence alignment between Homo sapiens CUL1, Saccharomyces cerevisiae CDC53, C. elegans CUL-1 and the C. elegans N2 and CB4856 allelic variants of CUL-6. The location of the N2 and CB4856 polymorphism is indicated with an arrow. The amino acid conservation is indicated by the grey bars at the bottom and by the annotations on top (single dot: weakly conserved, double dot: strongly conserved, asterisk: completely conserved). The gene cul-6 contains a polymorphism between N2 and CB4856 at a conserved site. Colors are based on the amino acid properties and locations of alpha-helices are indicated by cylinders [50]. C) Viral susceptibility of N2, CB4856, PHX1169 (N2 genetic background carrying a CB4856 cul-6 allele) and PHX1170 (CB4856 genetic background carrying a N2 allele) (asterisk indicating p <0.05, t-test).

To test whether the *cul-6* polymorphism explains the difference in viral susceptibility between N2 and CB4856 we used CRISPR-Cas9-modified strains encoding the *cul-6* N2 allele in the CB4856 genetic background (PHX1170) and the *cul-6* CB4856 allele in the N2 genetic background (PHX1169). Based on the results of the IL analysis PHX1170 was expected to be as susceptible as N2. We indeed observed that PHX1170 had a more susceptible phenotype, with a viral load in between that of N2 (p=0.41) and CB4856 (p=0.31) (Figure S3). PHX1169 retained high viral susceptibility with more variance in observed viral loads than in the N2 strain. Taken together, the *cul-6* polymorphism contributes to different viral susceptibility between N2 and CB4856, yet the effect size of this allele is modest. The resistant phenotype of CB4856 cannot be fully allocated to this allele, because it does not confer resistance in a susceptible background and CB4856 and PHX1170 were more similar than CB4856 and N2. This result shows that having the susceptible *cul-6* allele from N2 makes the strains vulnerable to infection while having the resistant allele from CB4856 does not protect strains with an otherwise susceptible N2 background.

## Discussion

Here we have unraveled the genetic architecture of viral susceptibility in the *C. elegans* strains N2 and CB4856. We found two QTL peaks on chromosome IV linking to susceptibility differences and confirmed the QTL on the right side of chromosome IV using a selection of ILs. Observations made for individual ILs show that multiple loci on chromosome IV contribute to viral susceptibility. When we zoomed in on the 12.4-12.9Mb region that likely contains a causal gene, we identified 34 polymorphic genes which may explain differences in viral susceptibility between N2 and CB4856. Allele swap experiments between one of these candidate genes, the IPR gene *cul-6*, indicated a single nucleotide polymorphism underlies susceptibility differences. Furthermore, we found that other genetic loci on chromosome IV also contribute to the whole phenotypic variation between N2 and CB4856. These findings show that the genetic architecture of OrV susceptibility is a complex, polygenic trait and future studies may identify even more genetic variants involved in OrV susceptibility.

### Chromosome IV is implicated in natural variation in OrV infection

By exposing RILs and ILs to OrV, we identified a QTL on chromosome IV that is implicated in a lower viral load due to the CB4856 allele. A genome wide association study (GWAS) on OrV infection in *C. elegans* also involved chromosome IV [15], but unlike this study, we did not find a QTL near the *drh-1* locus. This was in line with expectations as only two polymorphisms are found in the introns between N2 and CB4856 for this gene [27]. Still, the more distal associations uncovered by the GWAS could potentially result from the same allelic variation as the QTL between 12.41-12.89Mb, because the GWAS identified five locations on chromosome IV which are between 5 and 13Mb. Therefore, natural populations of *C. elegans* may carry similar genetic variants conferring OrV resistance as N2 and CB4856.

In our previous study investigating viral susceptibility differences between N2 and CB4856 we found that CB4856 had higher basal expression of IPR genes which we hypothesized may be caused by distinctive *pals-22*/*pals-25* expression patterns [26]. These genes, the respective repressor and activator of the IPR, are located adjacent to each other on the left of chromosome III [21]. eQTL studies showed local genetic variation (*cis*-eQTL) regulates expression of *pals-22* and *pals-25* [17,29,35,51–54]. Nevertheless, we did not observe a link between natural genetic variation in viral susceptibility in N2 and CB4856 and the *pals-22*/*pals-25* locus on chromosome III. Our results show that we could only explain a minor fraction of the heritability by the QTL locations we found. This result is typical for QTL mappings of complex traits and suggests that additional loci contribute to the viral susceptibility difference between N2 and CB4856. These loci may have small effect sizes, interactions or are affected by a (currently unknown) environmental cause [55].

### Orsay virus susceptibility has a polygenic basis

The QTL in the RIL panel and follow-up fine-mapping in the ILs identified a relatively small locus containing 34 polymorphic genes that contributes to the viral susceptibility towards OrV infection. We investigated the effect of a *cul-6* polymorphism and found that this SNP contributes to viral susceptibility. This allele functions in one direction by making the resistant CB4856 background susceptible when carrying the N2-allele. Yet, the phenotypic difference between N2 and CB4856 cannot be entirely explained by the *cul-6* allele alone. The 12.4-12.9Mb region also contains other genes that may affect viral susceptibility. Some of these are transcriptionally activated by OrV infection, others have more general or unknown cellular functions [16,19,22,24,49,56]. Besides, the 12.4-12.9Mb region specifically investigated here, we show that are multiple other loci and genes on chromosome IV contribute to viral susceptibility. The left side of chromosome IV appeared to be involved in determining the success of infection, but we could not verify this result in all the ILs. Studying this QTL may be complicated because only a small fraction of infections fails. Nevertheless, some ILs covering the left side of chromosome IV had a viral susceptibility distinct from the parent. Additionally, strain WN351 carries a susceptible introgression at the 12.4-12.9Mb locus but remained resistant. This strain has a large introgression also covering the left side of chromosome IV, where interacting genes may be located.

Our results reveal part of the complex genetic basis of OrV susceptibility. These results are in line with other studies mapping variation in viral susceptibility to the hosts’ genome (see for example [4,57–60]). Because viruses use the host’s cellular machinery to replicate and hosts have multiple mechanisms to counteract viruses, host-virus interactions will comprise many genetic interactions that can be affected by genetic variation. Thus, future studies may aim to uncover genetic networks rather than a single gene to further enhance our understanding of natural variation in host-virus interactions.

## Acknowledgements

The authors want to thank all the people that contributed to the OrV research in N2 and CB4856: Kobus Bosman, Henrikje Smits, Jikke Daamen, Koen Semeijn, Yahya Zakaria Abdou Gaafar, Maarten Costerus, Yuqing Huang, Emma Lagae and Niels Vissers. We want to thank Marie-Anne Félix for providing us with the OrV. We thank Daniel Cook, Robyn Tanny, and Erik Andersen for help in sequencing IL_CB4856_ strains. Lisa van Sluijs was funded by NWO (824.15.006) and Mark Sterken was supported by NWO domain Applied and Engineering Sciences VENI grant (17282).

## Author contributions

Conceived and designed the experiments: MGS, LvS, LBS, GPP, and JEK. Performed the experiments: MGS, LvS, YW, WR, MG, RV, and JAGR. Analyzed the data: MGS and LvS. Wrote the paper: MGS, LvS, GPP, and JEK, with contributions from all co-authors.

## Supplementary information

**Figure S1 Viral infection in L1 and L2 staged N2 and CB4856 –** A) Viral infections were either started in the L1 stage (22h) or L2 stage (28h). Populations were exposed to the OrV in liquid and isolated 30 hours post infection for viral load measurements. B) Viral loads observed for N2 and CB4856 that were infected in the L1 or L2 stage. Genotype had a significant effect on viral susceptibility for both stages (asterisk indicating p <0.05, t-test), but viral susceptibility was not determined by infection age.

**Figure S2 Genetic maps of the RIL and IL strains –** A) Genetic map of the N2xCB4856 RIL panel. B) Genetic map of chromosome IV for the N2-in-CB4856 background (strain WN340-WN354) and CB4856-in-N2 background (WN245-WN263) ILs.

**Figure S3 QTL mapping for mean, median and minimum viral load –** A) The QTL profile for mean viral load (including unsuccessful infections). The significant QTL peak is found from 12.5-14.9Mb. B) The QTL profile for median viral load (excluding unsuccessful infections). The significant QTL peak is found from 11.2-15.1Mb. C) The QTL profile for minimum viral load (including unsuccessful infections). The significant QTL peaks are found from 2.6-2.8Mb and 11.2Mb.

**Supplementary Table S1 IL strains used in this study –** Strain names, previously used strain names (in other publications), strain type, genotype and previous publications mentioning the strains are given.

**Supplementary Table S2 Genetic map of IL strains and markers** – A) Strain genetic data used to perform genetic linkage mappings. A reduced number of columns is shown for the strain map: the full-size table contains 384 columns. B) Informative marker positions used to perform genetic linkage mappings.

**Supplementary Table S3 Polymorphic genes between N2 and CB4856 at 12.41-12.89Mb on chromosome IV –** Selection of genes that have a polymorphism between N2 and CB4856 at the fine mapped QTL location. Gene name, location, strand, type of polymorphism, function according to WormBase and description in OrV literature are mentioned (format: first author, journal, year).

**Supplementary Text S1 Description of allele swap strains PHX1169 and PHX1170 –** Genetic code of the strains PHX1169 and PHX1170 compared to their respective parental genotypes N2 and CB4856

