## Supplementary Text S1 for "Punctuated loci on chromosome IV determine natural variation in Orsay virus susceptibility of *Caenorhabditis elegans* strains Bristol N2 and Hawaiian CB4856"

Strain name and genotype：PHX1169 cul-6(syb1169)

Synonymous mutation is labelled in blue.

PCR primers and sequencing primers:

SL01-Seq-s:AAGTGTTGTCTCTGAGTTGC

SL01-Seq-a:CGGATTAAGAGATCCTACGA

>SL01-syb1169

aagtgttgtctctgagttgctcatccatactatgtatgtactgccgtacaacttgggatccctgaataatttttcggaaaactactagaaacttccggaaagtttgttaaaactgtcagaaatgttgagcaaacttttttaaaagtttcagaaattttcagagttgttcaaaagtattgaattgaataccatttttagaaaaataaacattctaattctaaacaaacgaagtttcagGGTGTATCTCGAAAAGCGAATTTTAAAAGAAGGACACGAAACATTGCAAAGACTGGCTAAAGACAGTGGACTAAAAACTACACCAAAGGAATACATAACCAAACTCCTTGAAGTTCACGAGATTTATTTCAACTTGATAAATAAAGCATTCGATAGAAATGCACTTTTCATGCAATCTCTCGACAAAGCTTCTAAGGACTTTATCGAAGCTAATGCTGTGACTATGTTGGCTCCTGAAAAACACCGAAGCACAAGGTCTGCAGACTATCTTGCAAGATATTGTGATCAGCTCTTGAAGAAGAATTCC**AAA**GTTCAAGATGAAACGGCGCTAGATAAAGCAgtgagtttgcttttttcaactctgaacgtcgctgctccatcacatttttctgtacgaaaaattccctttatagatcaaaacatgatgggacagcctaacaccacgtatgtaaaaaaatcccattttcag**CTG**ACGGTGCTCAAG**TAC**ATCAGCGAAAAGGATGTCTTCCAATTGTATTATCAAAATTGGTTC**TCGAAGCGC**ATCATCAATAATTCTTCAGCAAGTGATGATGCAGAAGAAAAGTTCATTACAAATCTAACTGCAACAGAAGGTCTCGAATATACTCGCAATCTTGTGAAAATGGTCGAAGACGCTAAAATTAGTAAAGATTTGACTACTGAATTCAAAGATATCAAAACGGAAAAGTCGATTGACTTTAACGTTATTCTTCAAACAACTGGTGCATGGCCAAGTTTAGAT**CAG**ATCAAGATTATTCTTCCG**CGT**GAATTATCGACAATTCTCAAAGAGTTCGATACATTCTACAATGCGAGTCATAATGGACGAAGATTGAACTGGGCTTACTCACAATGCCGGGGCGAAGTTAACTCTAAAGCTTTCGAAAAGAAATATGTATTTATTgtgagtttatctatagttattcaaacttattatttttatcagataacattttattaaaaattttaaaatcagaagtcagcctgaagcagaagcagaagaagaagccaaactgctttttgaaatcttaacttattaaaagttagaattttccagGTTACAGCAAGCCAACTCTGCACGCTCTATCTTTTCAATGAGCAGGACTCGTTCACAATTGAACAAATTTCCAAAGCTATTGAAATGACTGCAAAGTCAACTTCGGCTATCGTAGGATCTCTTAATCCG

>wild type

aagtgttgtctctgagttgctcatccatactatgtatgtactgccgtacaacttgggatccctgaataatttttcggaaaactactagaaacttccggaaagtttgttaaaactgtcagaaatgttgagcaaacttttttaaaagtttcagaaattttcagagttgttcaaaagtattgaattgaataccatttttagaaaaataaacattctaattctaaacaaacgaagtttcagGGTGTATCTCGAAAAGCGAATTTTAAAAGAAGGACACGAAACATTGCAAAGACTGGCTAAAGACAGTGGACTAAAAACTACACCAAAGGAATACATAACCAAACTCCTTGAAGTTCACGAGATTTATTTCAACTTGATAAATAAAGCATTCGATAGAAATGCACTTTTCATGCAATCTCTCGACAAAGCTTCTAAGGACTTTATCGAAGCTAATGCTGTGACTATGTTGGCTCCTGAAAAACACCGAAGCACAAGGTCTGCAGACTATCTTGCAAGATATTGTGATCAGCTCTTGAAGAAGAATTCC**AAG**GTTCAAGATGAAACGGCGCTAGATAAAGCAgtgagtttgcttttttcaactctgaacgtcgctgctccatcacatttttctgtacgaaaaattccctttatagatcaaaacatgatgggacagcctaacaccacgtatgtaaaaaaatcccattttcag**TTA**ACGGTGCTCAAG**TAT**ATCAGCGAAAAGGATGTCTTCCAATTGTATTATCAAAATTGGTTC**AGTGAACGA**ATCATCAATAATTCTTCAGCAAGTGATGATGCAGAAGAAAAGTTCATTACAAATCTAACTGCAACAGAAGGTCTCGAATATACTCGCAATCTTGTGAAAATGGTCGAAGACGCTAAAATTAGTAAAGATTTGACTACTGAATTCAAAGATATCAAAACGGAAAAGTCGATTGACTTTAACGTTATTCTTCAAACAACTGGTGCATGGCCAAGTTTAGAT**CAA**ATCAAGATTATTCTTCCG**CGA**GAATTATCGACAATTCTCAAAGAGTTCGATACATTCTACAATGCGAGTCATAATGGACGAAGATTGAACTGGGCTTACTCACAATGCCGGGGCGAAGTTAACTCTAAAGCTTTCGAAAAGAAATATGTATTTATTgtgagtttatctatagttattcaaacttattatttttatcagataacattttattaaaaattttaaaatcagaagtcagcctgaagcagaagcagaagaagaagccaaactgctttttgaaatcttaacttattaaaagttagaattttccagGTTACAGCAAGCCAACTCTGCACGCTCTATCTTTTCAATGAGCAGGACTCGTTCACAATTGAACAAATTTCCAAAGCTATTGAAATGACTGCAAAGTCAACTTCGGCTATCGTAGGATCTCTTAATCCG

Strain name and genotype：PHX1170 cul-6(syb1170)

Synonymous mutation is labelled in blue.

PCR and sequencing primers:

SL02-SEQ-S:AAGTGTTGTCTCTGAGTTGC

SL02-SEQ-A:CGGATTAAGAGATCCTACGA

>SL02-syb1170

aagtgttgtctctgagttgctcatccatactatgtatgtactgccgtacaacttgggatccctgaataatttttcggaaaactactagaaacttccggaaagtttgttaaaactgtcagaaatgttgagcaaacttttttaaaagtttcagaaattttcagagttgttcaaaagtattgaattgaataccatttttagaaaaataaacattctaattctaaacaaacgaagtttcagGGTGTATCTCGAAAAGCGAATTTTAAAAGAAGGACACGAAACATTGCAAAGACTGGCTAAAGACAGTGGACTAAAAACTACACCAAAGGAATACATAACCAAACTCCTTGAAGTTCACGAGATTTATTTCAACTTGATAAATAAAGCATTCGATAGAAATGCACTTTTCATGCAATCTCTCGACAAAGCTTCTAAGGACTTTATCGAAGCTAATGCTGTGACTATGTTGGCTCCTGAAAAACACCGAAGCACAAGGTCTGCAGACTATCTTGCAAGATATTGTGATCAGCTCTTGAAGAAGAATTCC**AAA**GTTCAAGATGAAACGGCGCTAGATAAAGCAgtgagtttgcttttttcaactctgaacgtcgctgctccatcacatttttctgtacgaaaaattccctttatagatcaaaacatgatgggacagcctaacaccacgtatgtaaaaaaatcccattttcag**CTG**ACGGTGCTCAAG**TAC**ATCAGCGAAAAGGATGTCTTCCAATTGTATTATCAAAATTGGTTC**TCGGAGCGC**ATCATCAATAATTCTTCAGCAAGTGATGATGCAGAAGAAAAGTTCATTACAAATCTAACTGCAACAGAAGGTCTCGAATATACTCGCAATCTTGTGAAAATGGTCGAAGACGCTAAAATTAGTAAAGATTTGACTACTGAATTCAAAGATATCAAAACGGAAAAGTCGATTGACTTTAACGTTATTCTTCAAACAACTGGTGCATGGCCAAGTTTAGAT**CAG**ATCAAGATTATTCTTCCG**CGT**GAATTATCGACAATTCTCAAAGAGTTCGATACATTCTACAATGCGAGTCATAATGGACGAAGATTGAACTGGGCTTACTCACAATGCCGGGGCGAAGTTAACTCTAAAGCTTTCGAAAAGAAATATGTATTTATTgtgagtttatctatagttattcaaacttattatttttatcagataacattttattaaaaattttaaaatcagaagtcagcctgaagcagaagcagaagaagaagccaaactgctttttgaaatcttaacttattaaaagttagaattttccagGTTACAGCAAGCCAACTCTGCACGCTCTATCTTTTCAATGAGCAGGACTCGTTCACAATTGAACAAATTTCCAAAGCTATTGAAATGACTGCAAAGTCAACTTCGGCTATCGTAGGATCTCTTAATCCG

>CB4856

aagtgttgtctctgagttgctcatccatactatgtatgtactgccgtacaacttgggatccctgaataatttttcggaaaactactagaaacttccggaaagtttgttaaaactgtcagaaatgttgagcaaacttttttaaaagtttcagaaattttcagagttgttcaaaagtattgaattgaataccatttttagaaaaataaacattctaattctaaacaaacgaagtttcagGGTGTATCTCGAAAAGCGAATTTTAAAAGAAGGACACGAAACATTGCAAAGACTGGCTAAAGACAGTGGACTAAAAACTACACCAAAGGAATACATAACCAAACTCCTTGAAGTTCACGAGATTTATTTCAACTTGATAAATAAAGCATTCGATAGAAATGCACTTTTCATGCAATCTCTCGACAAAGCTTCTAAGGACTTTATCGAAGCTAATGCTGTGACTATGTTGGCTCCTGAAAAACACCGAAGCACAAGGTCTGCAGACTATCTTGCAAGATATTGTGATCAGCTCTTGAAGAAGAATTCC**AAG**GTTCAAGATGAAACGGCGCTAGATAAAGCAgtgagtttgcttttttcaactctgaacgtcgctgctccatcacatttttctgtacgaaaaattccctttatagatcaaaacatgatgggacagcctaacaccacgtatgtaaaaaaatcccattttcag**TTA**ACGGTGCTCAAG**TAT**ATCAGCGAAAAGGATGTCTTCCAATTGTATTATCAAAATTGGTTC**AGTAAACGA**ATCATCAATAATTCTTCAGCAAGTGATGATGCAGAAGAAAAGTTCATTACAAATCTAACTGCAACAGAAGGTCTCGAATATACTCGCAATCTTGTGAAAATGGTCGAAGACGCTAAAATTAGTAAAGATTTGACTACTGAATTCAAAGATATCAAAACGGAAAAGTCGATTGACTTTAACGTTATTCTTCAAACAACTGGTGCATGGCCAAGTTTAGAT**CAA**ATCAAGATTATTCTTCCG**CGA**GAATTATCGACAATTCTCAAAGAGTTCGATACATTCTACAATGCGAGTCATAATGGACGAAGATTGAACTGGGCTTACTCACAATGCCGGGGCGAAGTTAACTCTAAAGCTTTCGAAAAGAAATATGTATTTATTgtgagtttatctatagttattcaaacttattatttttatcagataacattttattaaaaattttaaaatcagaagtcagcctgaagcagaagcagaagaagaagccaaactgctttttgaaatcttaacttattaaaagttagaattttccagGTTACAGCAAGCCAACTCTGCACGCTCTATCTTTTCAATGAGCAGGACTCGTTCACAATTGAACAAATTTCCAAAGCTATTGAAATGACTGCAAAGTCAACTTCGGCTATCGTAGGATCTCTTAATCCG
